# Assessment of sensorimotor cortical beta oscillations from peripheral electromyography and force recordings

**DOI:** 10.64898/2026.07.07.736915

**Authors:** Christian Georgiev, Pierre Cabaraux, Nicolás Yanguma Muñoz, Daria Digileva, Scott J. Mongold, Vincent Wens, Armin Hakkak Moghadam Torbati, Lousin Moumdjian, Gilles Naeije, Xavier De Tiége, Mathieu Bourguignon

**Author notes:** These authors contributed equally to this work (Pierre Cabaraux, Nicolás Yanguma Muñoz). Corresponding Authors: Christian Georgiev and Mathieu Bourguignon, Laboratory of Functional Anatomy, Faculty of Human Motor Sciences, Université libre de Bruxelles (ULB), 1070 Brussels, Belgium.

## Abstract

The beta oscillations of the human primary sensorimotor cortex (SM1) play a crucial role in regulating motor and cognitive behavior in health and disease. However, their assessment relies on costly and complex neuroimaging techniques, limiting scalability and translational applications. We present a novel method for assessing beta oscillations from easily obtained peripheral electromyography and force recordings. We show that movement-induced modulations in SM1 beta oscillations can be assessed from the electromyography or force recordings of a contracted contralateral hand muscle. We demonstrate the fidelity of this method in young and elderly healthy participants and in Parkinson’s disease patients. We also demonstrate that the resting-state SM1 beta interhemispheric coupling can be assessed from an interhand coupling between the electromyography or force of contracted homologous hand muscles. This methodology enables scalable and cost-effective investigations of beta oscillations for all fields of human neuroscience and for the development of accessible disease/therapeutic markers.

## Main

### Oscillatory activity is a ubiquitous property of human cortical neural assemblies.^1,2^

In the primary sensorimotor cortex (SM1), beta oscillations at 13–30 Hz play a key role for motor control.^3^ Abnormalities in these oscillations are present in multiple disorders of movement execution and comprehension, such as Parkinson’s disease,^4^ stroke,^5^ autism,^6^ and schizophrenia.^7,8^ Thus, monitoring these oscillations allows for addressing fundamental questions in human neuroscience and could support diagnosis and rehabilitation. At present, non-invasive investigations of beta oscillations are only possible with electroencephalography (EEG) or magnetoencephalography (MEG). However, these methods are costly and require specialized infrastructure and expertise. More accessible measures of sensorimotor beta oscillations would transform the ability to conduct large-scale translational research.

Electrophysiological recordings from the SM1 are characterized by bursty 13–30 Hz beta oscillations.^9–13^ A suppression of these beta bursts, resulting in suppressed global beta amplitude, is observed during movement preparation and execution and is typically followed by a rebound after movement termination.^14–17^ Hence, the beta oscillations are an indicator of the state of activation of the SM1, with amplitude suppression, termed event-related desynchronization (ERD), indicating SM1 excitation and amplitude enhancement, termed event-related synchronization (ERS), indicating SM1 inhibition.^18,19^ The beta ERD and ERS typically begin in the active SM1 and become bilateral shortly after, resulting in a coupling between the left and right SM1s.^13,16^ This interhemispheric coupling is also present at rest and defines a bilateral network associated with motor performance.^20,21^

Several findings suggest that beta oscillations could be reliably assessed from cost-effective recordings from beyond the cortex. Beta oscillations define periods of excitation and inhibition for SM1 pyramidal neurons. Hence, the corticospinal drive tends to couple in phase and amplitude with these oscillations.^22^ Thus, SM1 beta oscillations propagate to spinal alpha motor neurons,^23^ and shape the activity of skeletal muscles.^24–26^ This is attested by the existence of beta-band coupling between SM1 activity and electromyographic (EMG) activity of isometrically contracted muscles, known as corticomuscular coherence (CMC).^22,27–29^ In addition, a coupling similar to CMC has been reported between SM1 activity and contraction force.^27^ Therefore, beta oscillations also propagate to the force output of isometrically contracted muscles.

Here, we investigate the feasibility of assessing the movement-related SM1 beta ERD from peripheral (EMG or force) recordings. For that, we introduce tasks that combine a steady muscle contraction with transient movements of another body segment. In that setup, the movement is expected to modulate SM1 beta oscillations and we hypothesize that a trace of this modulation can be recorded from the EMG or force of the steadily contracting muscles. Additionally, we hypothesize that the resting-state SM1 beta interhemispheric coupling leads to a similar interhand coupling between EMGs or forces of contracted homologous bilateral hand muscle. Validating this hypothesis would extend the method we present to connectivity analyses. Finally, we assess the validity of the developed method by replicating previous findings of age-related increase in SM1 beta ERD^30–32^ and by determining its applicability to the challenging experimentation with Parkinson’s disease patients.

## Results

### Movement-Related SM1 and Peripheral Beta ERD

We first introduce a dual-movement task that enables the characterization of movement-induced beta amplitude modulation in peripheral (EMG and force) signals and investigate the extent to which the peripheral modulations are a reflection of the SM1 modulation recorded with MEG. Fourteen healthy young adults (group 1) held an isometric pinch grip contraction with their right hand and performed slow transient rotational movements of the right brachium in response to auditory cues presented every 3–4 s (Fig. 1a–b). The brachium movement leads to a large beta modulation in the SM1 area governing that muscle group that spreads across the SM1 and reaches neighboring SM1 areas,^33^ including the one governing the isometrically contracted right first dorsal interosseous (FDI). Therefore, the isometric contraction serves as a probe to collect the beta oscillations and their modulation induced by the brachium movement. MEG, surface EMG from the FDI muscle, the applied contraction force, and the acceleration of the right hand were simultaneously recorded. MEG, EMG, and force signals were subjected to a time-frequency analysis from -1 to 2.5 s relative to brachium movement onset as detected via accelerometry. In each signal, beta ERD was quantified in terms of size and depth.

**Fig. 1.**
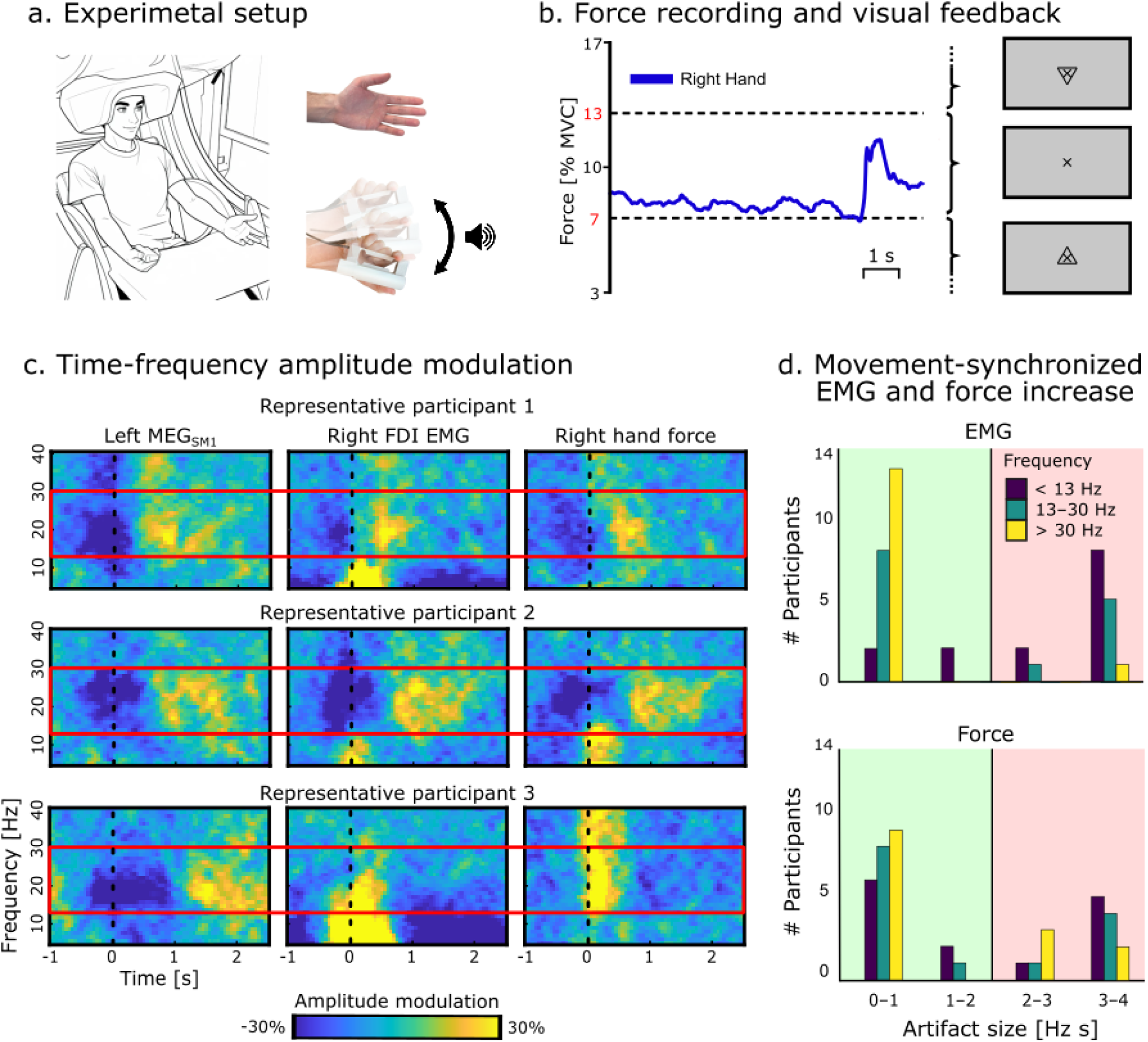
Ipsilateral upper limb dual contraction–movement execution task. **a.** Experimental setup. Participants held an isometric pinch grip contraction with their right hand and performed slow transient rotational movements of the right brachium in response to auditory cues presented every 3–4 s. **b.** Force recording, expressed as % Maximum Voluntary Contraction (MVC), and visual feedback to ensure contraction stability. Arrowheads pointing up or down instructed the participants to increase or decrease the force when necessary. **c.** Time-frequency amplitude modulation of left MEG_SM1_, right FDI EMG, and right hand force signal for two representative participants who successfully performed the task (first 2 rows) and one representative participant who did not manage to sustain a stable force (3^rd^ row). Dotted black lines at time 0 indicate movement onset and red rectangles outline the classical beta frequency range (13–30 Hz). **d.** Histograms of the size of movement artifacts in FDI EMG and force maps at frequencies below, within, and beyond the beta range. Black vertical lines indicate the artifact significance threshold of 2 Hz s.

Performing the right brachium movement resulted in a large-scale beta ERD in the MEG signals above both SM1s (MEG_SM1_), along with a similar ERD in right EMG and force signals in the time interval from -0.5 to 1 s (Fig. 1c; Supplementary Table 1). Compared to the left MEG_SM1_, the ERD size was significantly smaller in the peripheral signals (EMG, *t*_13_ = 3.32, *p*_corrected_ = 0.018; force, *t*_13_ = 3.79, *p*_corrected_ = 0.007), although its depth did not differ significantly (EMG, *t*_13_ = 1.23, *p*_corrected_ = 0.72; force, *t*_13_ = 2.56, *p*_corrected_ = 0.072; Supplementary Table 1). An inspection of the peripheral signals revealed that the overall decrease in beta ERD size compared to left MEG_SM1_ was due to the inability of some participants to sustain a stable contraction throughout the task. In these cases, the execution of brachium movements tended to co-occur with an increase in EMG and force fluctuations which likely obscured the beta ERD (Fig. 1c bottom row). Accordingly, 6 participants had a significant amplitude increase around brachium movement onset in the EMG that reached the beta band and 5 participants had a similar artifact in the force (Fig. 1d). Of note, a larger proportion of participants displayed a low frequency (<13 Hz) movement-synchronized artifact in both peripheral signals, and fewer participants displayed such an artifact at frequencies beyond the beta band (>30 Hz), predominantly in the force signal (Fig. 1d). However, these artifacts are less problematic as they do not affect the detectability of the SM1 beta propagation effect.

To circumvent the issue of unstable EMG and force during the movement, we designed a different movement execution task which takes advantage of the well-established bilaterality of the movement-related SM1 beta ERD.^13,16,26^ Forty one participants (group 2) held an isometric pinch grip contraction with their right hand and performed a closing/opening movement of the left fingers in response to auditory cues presented every 3–4 s. In addition to a strong beta ERD in the right MEG_SM1_, we expected to observe a similar beta ERD in left MEG_SM1_ which propagated to the right EMG and force signals.

Performing the left finger movement resulted in a large-scale beta ERD in both MEG_SM1s_, along with a similar ERD in right EMG and force signals in the time interval from -0.5 to 1 s. Compared to left MEG_SM1_, the ERD size was significantly smaller in both the EMG (*t*_40_ = 6.96, *p*_corrected_ < 0.001) and the force (*t*_40_ = 6.76, *p*_corrected_ < 0.001), and its depth was also significantly smaller in EMG (*t*_40_ = 3.71, *p*_corrected_ = 0.001) and force (*t*_40_ = 6.60, *p*_corrected_ < 0.001; Supplementary Table 2). With this task, participants were more successful in maintaining a stable right hand contraction force (Fig. 2a–b) resulting in a better ability to detect beta oscillatory activity in the peripheral signals. Two participants displayed a significant movement-synchronized amplitude increase in the EMG beta band and 3 participants displayed such an artifact in the force (Fig. 2b; Supplementary Fig. 2). However, in one participant, what was identified as a force artifact was consistent with an early beta event-related synchronization (Supplementary Fig. 2).

**Fig. 2.**
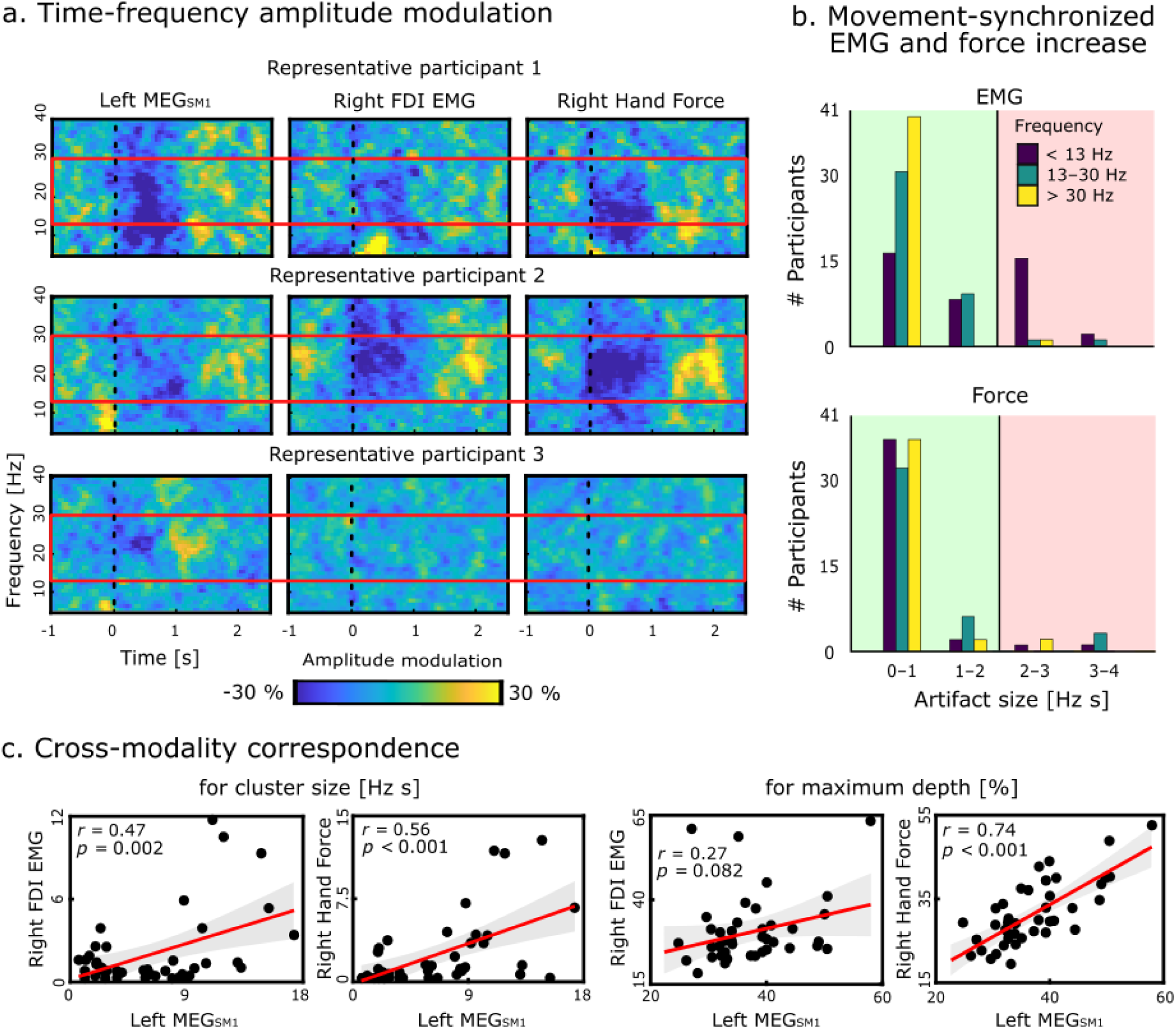
Contralateral upper limb movement execution task. **a.** Time-frequency amplitude modulation of left MEG_SM1_, right FDI EMG, and right hand force signals for three representative participants who successfully performed the task. Participants 1 and 2 displayed beta ERD for all signals, but participant 3 showed an ERD only for MEG_SM1_ signals. Dotted black lines at time 0 indicate movement onset and red rectangles outline the classical beta frequency range (13–30 Hz). **b.** Histograms of the size of movement artifacts in FDI EMG and force maps at frequencies below, within, and beyond the beta range. Black vertical lines indicate the artifact significance threshold of 2 Hz s. **c.** Cross-modality Pearson correlations for the size of the cluster of significant beta suppression (left) as well as its maximum depth (right). Regression lines are in red and associated 95% confidence areas are in gray.

A cross-modality correlation analysis revealed strong positive correlations between left MEG_SM1_ and peripheral signals for their beta ERD size and depth (Fig. 2c). Besides, correlation analyses between the time-frequency maps of the 3 signals in the 13*–*30 Hz and -0.5 to 1 s time-frequency window revealed positive correlations between the left MEG_SM1_ and peripheral signals (right EMG, *r* = 0.13 ± 0.35, *t*_40_ = 2.41, *p* = 0.021; force, *r* = 0.26 ± 0.32, *t*_40_ = 5.30, *p* < 0.001), indicating a moderate but significant similarity between the patterns of beta amplitude variation of the 3 signals.

Further analyses investigating beta ERS are presented in Supplementary text S1. Moreover, further analyses investigating the changes in the properties of the beta bursts underlying the ERD are presented in Supplementary text S2.

### Consistency of Beta ERD Across Tasks

To further validate the *contralateral dual movement execution task*, we quantified the bilaterality of beta ERD and compared its depth and size to those of the *ipsilateral dual movement task* for group 1 participants with artifact-free data. The results of the two tasks consistently revealed that MEG_SM1_ beta ERD was bilateral. In addition, left and right MEG_SM1_ beta ERD size and depth showed some degree of consistency across tasks (left size: *r* = 0.64, *p* = 0.013; right size: *r* = 0.48, *p* = 0.08; left depth: *r* = 0.80, *p* < 0.001; right depth: *r* = 0.40, *p* = 0.16), suggesting an idiosyncratic, yet stable, pattern of SM1 beta ERD for each participant. Furthermore, for participants who successfully maintained a stable contraction throughout both tasks, the patterns of peripheral beta ERD were mostly consistent across tasks as revealed by positive correlations across tasks for EMG ERD size (EMG, *r* = 0.85, *p* = 0.007) and force ERD depth (*r* = 0.77, *p* = 0.024). The correlations for force ERD size and EMG ERD depth were also strong across tasks, however, did not reach significance likely due to limited sample size (force ERD size, *r* = 0.66, *p* = 0.073; EMG ERD depth, *r* = 0.60, *p* = 0.11).

### Central and Peripheral Beta Interhemispheric Coupling

Given that beta oscillations propagate to peripheral signals during isometric contractions, the interhemispheric coupling between the two SM1s should be reflected by an interhand coupling between EMGs or forces. To test this hypothesis, 35 healthy young adults (group 3) held an isometric pinch grip contraction with both hands while MEG, surface EMG from both FDI muscles, and the applied force from both hands were simultaneously recorded (Fig. 3a–b). Correlations between the beta-band envelopes of source-reconstructed left and right SM1 were estimated to quantify beta interhemispheric coupling. Similar correlation analyses were applied between the left and right FDI EMGs and the left and right force signals.

**Fig. 3.**
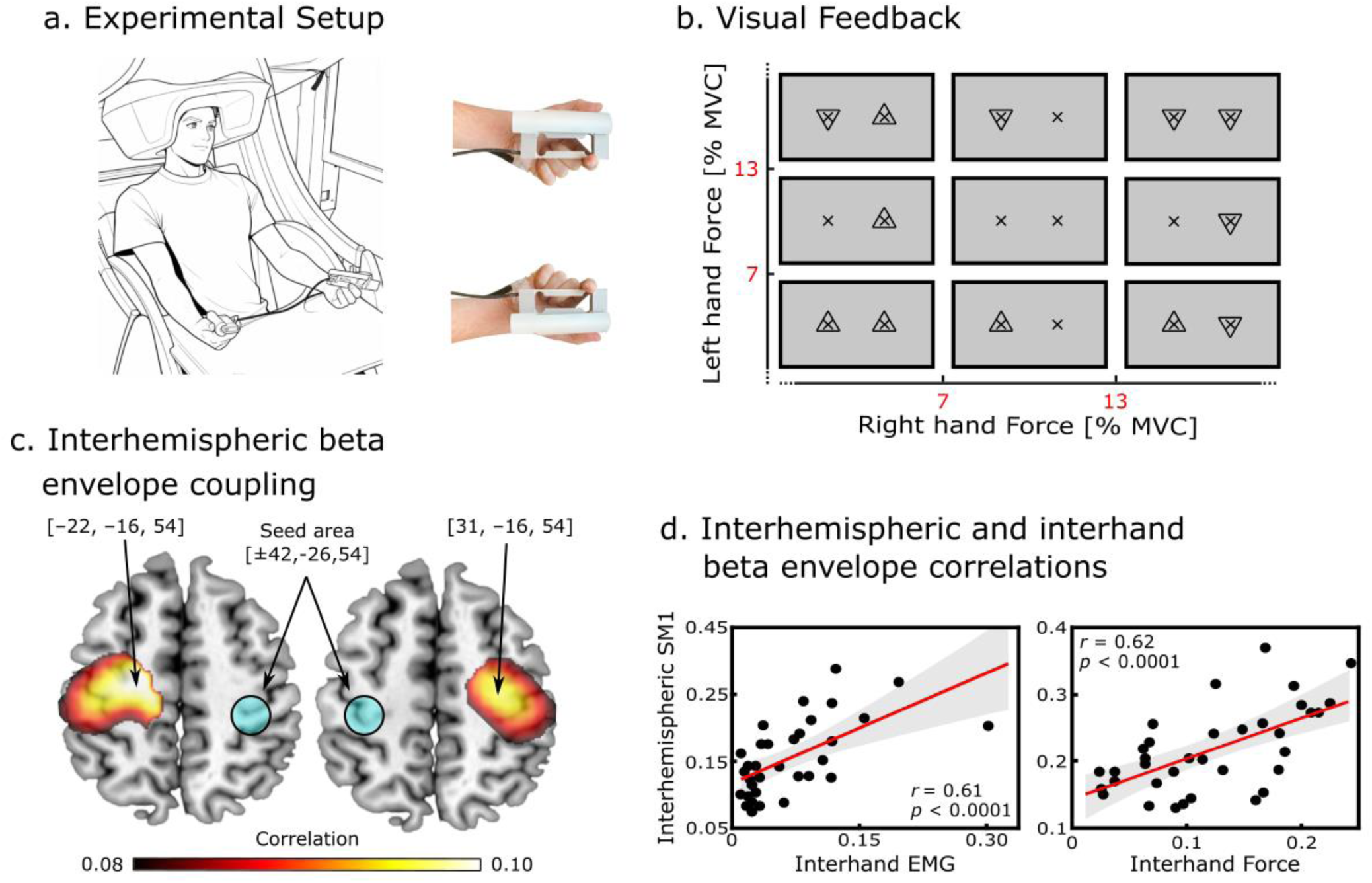
Bimanual contraction task. **a.** Experimental setup. Participants held an isometric pinch grip contraction with their both hands. **b.** Force visual feedback to ensure contraction stability. **c.** Maps of beta interhemispheric coupling between the left and right SM1 hand regions. To limit computational cost, maps for a given seed area were estimated only for sources within a 4 cm sphere centered on the corresponding area in the opposite hemisphere. **d.** Pearson correlations between averaged SM1 interhemispheric beta envelope coupling and interhand EMG (left) and force (right) beta envelope couplings. Regression lines are in red and associated 95% confidence areas are in gray.

A two-seed-based connectivity analysis revealed that the beta envelope in the left SM1’s hand area (Montreal Neurological Institute (MNI) coordinates: [-42 -26 54] mm) was significantly correlated with homologous regions in the right SM1 (MNI coordinates: [30 -25 49] mm) and vice versa (Fig. 3c, Supplementary Fig. 3). Across participants, maximal coupling occurred at 16.8 ± 1.7 Hz frequencies.^34^

Similarly, there was a significant beta envelope correlation between the left and right FDI EMGs (*r* = 0.12 ± 0.06, *p* < 0.001), and between the left and right force signals (*r* = 0.07 ± 0.06, *p* < 0.001; Supplementary Fig. 3). Across participants, maximal interhand coupling occurred at 19.4 ± 3.0 Hz for the EMG and 18.7 ± 3.0 Hz for the force.

A cross-modality correlation analysis revealed a significant positive correlation between participants’ interhemispheric and interhand beta envelope couplings (SM1*–*EMG, *r* = 0.61, *p* < 0.0001; SM1*–*force, *r* = 0.62, *p* < 0.0001; Fig. 3d). There was also a strong association across participants between the frequency of peak interhemispheric and interhand couplings (SM1*–*EMG, *r =* 0.67, *p* = 0.012; SM1*–*force, *r* = 0.69, *p* = 0.004). Collectively, these positive associations suggest that the beta-band coupling between the left and right peripheral signals is a mirrored reflection of the beta-band coupling between the left and right SM1s.

Further analyses confirmed that interhemispheric and interhand beta envelope couplings originate from the presence of beta bursts in MEG and peripheral signals (Supplementary text S3).

### Validation of Peripheral Beta Assessment Methodology

We further validated the methodology for assessing beta oscillations from the periphery at two levels. First, we demonstrated its applicability outside the magnetically shielded room of the MEG and with cost-effective and easily accessible data acquisition equipment. Second, we demonstrated that it allows for the identification of group differences in ERD, illustrated by a comparison between young and older adults, the latter being known to display greater ERD.^30^ Twenty-one young (29 ± 5 years old) and 13 older (66 ± 6 years old) participants (group 4) performed the *contralateral finger movement execution task* in standard laboratory settings. The force of the right hand and the acceleration of the left hand were recorded and analyzed as previously described.

A significant beta ERD in the time interval from -0.5 to 1 s relative to left finger movement was identified in 12 (57%) young participants, and in 10 (77%) older participants in the force signal (all *p* < 0.05). This beta ERD was greater in older compared to young participants in terms of size (young, 1.85 ± 2.20 Hz s; older, 6.60 ± 5.20 Hz s; *t_32_* = -3.71, *p* < 0.001) and depth (young, 23 ± 7%; older, 34 ± 11%; *t_32_* = -3.34, *p* = 0.002; Supplementary Fig. 4). Further analyses investigating age differences in ERS are presented in Supplementary text S1.

### Peripheral Beta Assessment in Parkinson’s Disease

Finally, we assessed the feasibility of assessing beta oscillations from the periphery in a clinical population affected by resting tremor. Thirteen Parkinson’s disease patients (group 5; Supplementary Table 4) performed the contralateral finger movement execution task in standard laboratory settings in OFF medication condition. Beta ERD was monitored with EEG and force recordings.

A significant beta ERD in the time interval from -0.5 to 1 s was identified in 13 (100%) patients for both the left and right EEG_SM1_ and in 12 (92%) patients for the force (*p* < 0.05). However, in 2 patients, the force beta ERD was obscured by a broadband movement-synchronized amplitude increase. This amplitude increase stemmed from movement artifacts in 1 patient and was of unclear origin in the second patient (Fig. 4). A cross-modality correlation analysis identified a negative correlation between EEG_SM1_ and force time-frequency amplitude modulation maps for these 2 patients (*r* = -0.39 and -0.65, respectively). In the remaining 11 patients, the cross-modality correlation was positive (*r* = 0.49 ± 0.25, *t*_10_ = 6.5, *p* < 0.001). Moreover, the cross-modality correlation was significantly positive across all patients (*r* = 0.34 ± 0.45, *t*_12_ = 2.7, *p* = 0.019). EEG_SM1_ and force beta ERD cluster size and depth are presented in Supplementary Table 5. Further analyses investigating the beta ERS in the EEG_SM1_ and the force signals can be found in the Supplementary text S1 and in Supplementary Table 5.

**Fig. 4.**
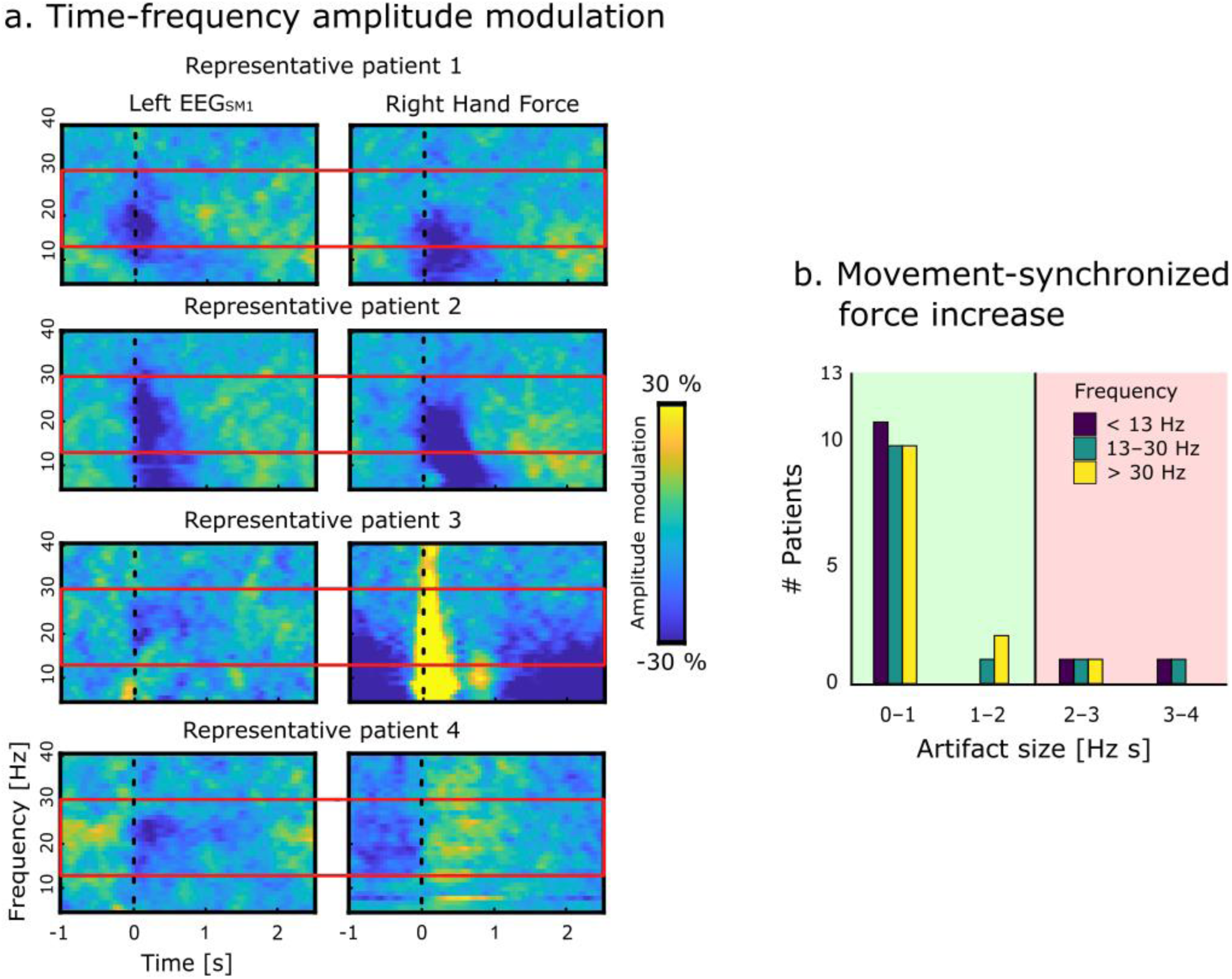
Beta modulations in Parkinson’s disease patients executing the contralateral upper limb movement execution task. **a.** Time-frequency amplitude modulation in the EEG_SM1_ and force signals of 4 Parkinson’s disease patients. The first two rows depict patients in whom a beta ERD was clearly seen in both the left EEG_SM1_ and the right hand force. The third and fourth rows depict the two patients for whom a beta ERD was clearly visible in the left EEG_SM1_, and obscured by movement-synchronized artifacts in the right hand force. Dotted black lines at time 0 indicate movement onset and red rectangles outline the classical beta frequency range (13–30 Hz). **b.** Histograms of the size of movement artifacts in force maps at frequencies below, within, and beyond the beta range. Black vertical lines indicate the artifact significance threshold of 2 Hz s.

## Discussion

The beta oscillations reflect the state of activation of the SM1s.^12,16,35,36^ Their assessment is ubiquitous in basic and clinical neuroscience.^30,36^ We introduce a novel approach to measure the SM1 beta oscillations using cost effective and easily applicable EMG and force recordings. Building on the knowledge that SM1 beta oscillations propagate to skeletal muscles during steady muscle contraction, we developed tasks that combine a steady muscle contraction with transient movements of another body segment. In that setup, the movement triggers bilateral modulations in SM1 beta oscillations, a trace of which can be recorded from the EMG or force of the steadily contracting muscle. This approach was also extended to assess beta interhemispheric coupling by evaluating interhand beta envelope coupling during bilateral steady muscle contraction.

The movement execution induced a bilateral SM1 beta ERD.^14–17^ Similar beta ERD could be recorded from the EMG and the force of a contracted hand muscle. The peripheral ERD was correlated with the SM1 ERD, suggesting a good correspondence between the two. Importantly, we investigated movement executions either by the upper limb that maintains the steady contraction or by the contralateral one. The ipsilateral task was more challenging, resulting in unstable EMG and force, thereby obscuring the true beta propagation effect. Therefore, the contralateral task appears better suited for this purpose, by leveraging the well-documented bilaterality of movement-induced SM1 beta ERD.^26, 37^

We identified a strong correlation between the SM1 beta envelope interhemispheric coupling and analogous interhand EMG or force coupling during bilateral contraction, indicating that the latter are a valid proxy of the former. This expands the scope of our method to the realm of functional connectivity analyses and opens perspectives for more easily obtained assessments of the sensorimotor network.

Further validating the proposed approach, we show that beta ERD can be extracted from a contraction force signal outside a magnetically shielded room and with very affordable data acquisition equipment and freely available software. In addition, using beta ERD collected from force signals, we could replicate the well-documented increase in ERD amplitude with aging.^30–32^ Finally, we demonstrate that the proposed approach to extract beta ERD from force signals, especially with the *contralateral dual-movement execution task*, can be applied to most individuals with Parkinson’s disease despite their motor impairments.

Some limitations to the proposed methods should be highlighted. First, the approach places participants in dual-task conditions. This could be an issue in conditions where cognition is altered, or for applications where attention should be focused on other tasks, such as watching videos to assess beta modulations in response to action observation. Second, the degree to which beta SM1 oscillations are seen in peripheral signals shows substantial inter-individual variability. This was highly expected given the known variability in the magnitude of CMC.^27,38^ However, most clinical applications will concern aging populations, where CMC is higher,^28^ and where beta ERD in the force was most salient in our data. Besides, the magnitude of scalp-detected SM1 oscillations also depends on individual factors, such as skull thickness and source orientation. This is, however, not an issue in applications where inter-individual variability is not of interest, or in longitudinal designs. And finally, in subjects with limited beta ERD in the force, experiment duration could be increased. In fact, upon continuous evaluation of time-frequency amplitude modulation, a stop criterion for the recordings could be the detection of a significant beta ERD.

Overall, our results support the reliability of a novel method for easily accessible and inexpensive recordings of the SM1 beta oscillations in the absence of neuroimaging techniques such as EEG or MEG. Although these techniques are the current gold standard in the field, our novel assessment method allows for accessible, yet equally resolute, recordings of the changes in beta oscillations which accompany motor behavior. Moreover, our method provides great advantages for recordings with special populations such as children or clinical populations, for whom the application of EEG or MEG may not be feasible. Therefore, it paves the way for unprecedented advances in the assessment of the diagnostically-relevant properties of beta oscillations in hospitals, nursing homes, rehabilitation centers, or even at home.

## Methods

### Participants

Fourteen healthy volunteers (group 1; 7 female; Mean age 27 ± 6 years) participated in the MEG *ipsilateral upper limb dual contraction-movement execution task*. Those 14 along with 27 additional healthy volunteers (group 2; 10 female; Mean age, 26 ± 3 years) participated in the MEG *contralateral upper limb dual contraction-movement execution task*. Thirty-five healthy volunteers (group 3; 13 female; Mean age, 26 ± 5 years) participated in the MEG *bimanual contraction task*. In addition, 34 healthy volunteers (group 4), of whom 21 were young (10 female; Mean age, 29 ± 5 years) and 13 were older (8 female; Mean age, 66 ± 6 years), as well as 13 patients with Parkinson’s disease (group 5; 7 female; Mean age, 67 ± 6 years; see Supplementary Table 4 for patients’ clinical characteristics) participated in validation recordings. All healthy participants and 9 of the Parkinson’s disease patients were right-handed as assessed by the Edinburgh Handedness Inventory,^39^ with no known neurological or psychiatric disorders (other than Parkinson’s disease in the patient group), and had normal or corrected-to-normal vision. The study was approved by the local ethics committee (Comité d’Ethique Hospitalo-Facultaire Erasme-ULB, 021/406; Reference: B4062021000049) and conducted in accordance with the Declaration of Helsinki. All participants gave a written informed consent prior to participating.

### Experimental Protocols

Three experimental tasks were used for this study: an *ipsilateral* and *contralateral upper limb dual-movement execution task*, and a *bimanual contraction task*. Prior to recording, each participant’s FDI maximum voluntary contraction (MVC) force was identified during a 5-second maximal strength pinch grip contraction against a 500-N force transducer (model CS 50-3Q1, SAUTER GmbH, Wutöschingen, Germany). During the recordings, participants of groups 1, 2, and 3 were seated inside a magnetically shielded room with their head inside a MEG helmet (Fig. 1a). Participants of groups 4 and 5 were seated in a chair in standard laboratory settings (outside the magnetically shielded room of the MEG) and, in the case of group 5, were wearing an EEG cap.

### Dual-movement execution tasks

Participants were asked to maintain a stable isometric contraction in the form of a pinch grip with their right thumb and index finger at 10 ± 3% MVC force for a duration of 5 minutes. For both tasks, the applied force was recorded with a custom-made 6-N force transducer (with a load cell model № 1004, Vishay Precision Group, Malvern, PA, USA equipped with a 3D printed handle). Participants were asked to fixate their gaze on a fixation cross presented on a MEG-compatible projection screen (groups 1 and 2) or on a 16-inch HD computer screen (groups 4 and 5) placed approximately 1 meter in front of them. Real-time visual feedback informed the participants when the contraction force was no longer within 10 ± 3% MVC (Fig. 1b), allowing for correction of their applied force. For the *ipsilateral upper limb dual-movement execution task*, while holding the isometric contraction with the right hand, an auditory cue (presented with random jitter every 3–4 s) prompted the participants to perform a 5–10 degree external or internal rotation of the right brachium alternately, which resulted in a small lateral movement of the contracting hand in the horizontal plane (Fig. 1a). To limit artifacts in peripheral signals, participants were instructed to complete the rotation at a slow pace, in about half a second. For the *contralateral upper limb dual-movement execution task*, while holding the isometric contraction with the right hand, an identical auditory cue prompted the participants to perform a fast closing/opening movement with the fingers of the left hand. Participants of group 1 performed the two tasks in a random order, while participants of group 2 performed the *contralateral upper limb dual-movement execution task* once and participants of groups 4 and 5 performed the same task twice.

### Bimanual contraction task

Participants of group 3 were comfortably seated with their head inside the MEG helmet and with both hands placed on the supporting table (Fig. 3a). For two 5-minute blocks, they were asked to maintain a steady bimanual isometric contraction in the form of a pinch grip with their thumb and index finger at 10 ± 3% of each hand’s MVC force. The force recordings and the real-time feedback procedure were identical to the ones used in the *dual-movement execution tasks*, this time with force feedback being provided for each of the two hands separately (Fig. 3b).

### Data acquisition

#### Magnetoencephalography (MEG; groups 1, 2, and 3)

All MEG recordings were conducted at the HUB Hôpital Erasme (Université Libre de Bruxelles, Brussels, Belgium) in a lightweight magnetically shielded room (Maxshield, Elekta, Helsinki, Finland) with a whole-head 306-channel neuromagnetometer system (TRIUX, MEGIN, Helsinki, Finland) that comprises 102 magnetometers and 102 planar gradiometer pairs. MEG signals were recorded using a bandpass of 0.1–330 Hz and a sampling rate of 1 kHz. Throughout the recording, head position was monitored by passing current to 4 head-tracking coils attached to each participant’s head. Prior to the acquisitions, the locations of the coils and at least 300 other head-surface points with respect to anatomical fiducials were digitized with an electromagnetic tracker (Fastrak, Polhemus, Colchester, VT, USA).

For source-space analyses of the MEG activity, anatomical 3D T1 scans were obtained for each participant with a 3 Tesla magnetic resonance scanner (Signa, General Electric, New York, NY, USA).

#### Electroencephalography (EEG; group 5)

EEG was recorded using a 64-channel EEG cap (EEGO Mylab, ANT Neuro, Hengelo, the Netherlands), with electrodes arranged in accordance with an augmented 10–20 system. Signals were sampled at 1 kHz. Electrode impedance was kept below 20 kΩ via the application of electroconductive gel and the recording was online referenced to electrode CPz.

#### Electromyography (EMG), force, and accelerometry recording

Monopolar surface EMG was recorded—for groups 1, 2, and 3 only—from the left and right FDI muscles with an active Ag–AgCl electrode on the muscle belly and a reference on the first metacarpal bone. Electroconductive gel was applied between the electrode surface and the skin. The force of the contracted FDI muscle was recorded with the 6N force transducer (see section ***Dual-movement execution tasks***). For the *dual-movement execution tasks*, hand movement acceleration was recorded with a MEG-compatible 3-axis accelerometer (ADXL335, AnalogDevices, Wilmington, MA, USA) attached either to the right wrist during the *ipsilateral upper limb movement execution task* or to the left index finger during the *contralateral upper limb movement execution task*. The accelerometer was always attached in a way that did not impede the participants’ movement.

For groups 1, 2, and 3, EMG, force, and accelerometer signals were acquired with the additional bipolar channels of the MEG system and hence recorded time-locked to the MEG signals with a passband of 10–330 Hz (EMG) or DC–330 Hz (force and acceleration). For groups 4 and 5, force and accelerometer signals were recorded in synchrony at 1 kHz with a multifunction MCC USB-231 data acquisition device (Measurement Computing Corporation, Newbury, United Kingdom) via the free DAQami™ 4.2.1 software.

### Data preprocessing

The MEG data from all tasks were preprocessed off-line with temporal signal-space separation (window length, 10; correlation limit, 0.9) for elimination of external signal sources and correction of head movement artifacts.^40^ Then, MEG signals were band-pass filtered through 0.3–45 Hz and subjected to independent components analysis^41^ for further artifact suppression. Thirty independent components were evaluated from the MEG data with a FastICA algorithm (dimension reduction, 30; nonlinearity, tanh).^41,42^ Between 3 and 5 components per participant were removed from the data, corresponding to ocular, myogenic, and cardiac artifacts.

The same filtering was applied to the EEG data from the Parkinson’s disease patients. Following a standard preprocessing pipeline,^43^ bad EEG channels were identified and topographically interpolated from the signals of neighboring channels and the EEG was rereferenced to a common average. Lastly, independent components analysis was also applied to the EEG data (dimension reduction, 20; nonlinearity, tanh). Between 2 and 6 components per participant were removed from the data, corresponding to ocular, myogenic, and cardiac artifacts.

Time points less than 1 s away from periods when MEG or EEG amplitude exceeded 5 standard deviations above the mean or when force visual feedback was present were considered contaminated by artifacts.

### Data Analyses

#### SM1 and Peripheral Beta amplitude modulation

For investigating the magnitude of beta ERD in the *movement execution tasks*, segments of artifact-free MEG, EEG, EMG, and force data were detrended and filtered through 5-Hz-wide frequency bands centered on 5–40 Hz by 1 Hz steps. The bandpass filters used for this were designed in the frequency domain with zero-phase and 1-Hz-wide squared-sine transitions from 0 to 1 and 1 to 0 (e.g., the filter centered on 20 Hz rose from 0 at 17 Hz to 1 at 18 Hz and ebbed from 1 at 22 Hz to 0 at 23 Hz). Further MEG analyses were conducted on gradiometer data, with each of the 102 gradiometer pairs within each subband combined using the first principal component of the signal pair. Subband signals’ envelope was extracted with the Hilbert transform and downsampled to 20 Hz. Segments were then trimmed by 1 s on each extremity to avoid filtering edge effects. Following, the subband envelopes were divided by their low-pass filtered version at 0.1 Hz (squared-sine transition from 0.05 to 0.15 Hz) to avoid contamination by very slow envelope variations. Such normalization ensures that envelopes fluctuate around 1, and are blind to changes slower than 0.1 Hz that cannot be ascribed to the task. Normalized envelopes were subsequently segmented into epochs of 3.5 s, from -1 to 2.5 s relative to either ipsilateral side brachium movement onset or contralateral finger movement onset. Movement onset was identified based on a threshold change in the signal of the accelerometer attached to the moving hand, and further corrected where necessary following visual inspection. As a result of the preprocessing steps, epochs which contained time points less than 2 s away from artifacts were not included in this analysis. On average, the number of retained epochs for the *ipsilateral upper limb movement execution task* was 63 (out of 85) in group 1. For the *contralateral upper limb movement execution task*, it was 67 (out of 85) in group 2, 142 (out of 170) in group 4, and 102 (out of 170) in the Parkinson’s disease patients of group 5. Normalized envelopes were averaged across epochs for each participant, giving rise to a time-frequency map of amplitude modulation for each MEG, EEG, EMG, or force channel. For MEG data, all subsequent analyses were performed on a preselection of 9 gradiometer pairs located above the left SM1 and its right hemisphere counterpart.^27,38^ For each SM1, we further retained the gradiometer pair which showed the maximal amplitude suppression in a preselected time-frequency window ranging from -500 ms before to 1500 ms after movement and from 13 to 30 Hz. The same approach was used for EEG analyses, where the left SM1 selection consisted of C1, C3, C5, CP1, CP3, CP5, FC1, FC3, FC5, and the right SM1 selection consisted of the right homologous electrodes. In 1 Parkinson’s patient the EEG was contaminated by large muscle artifacts. In this case, we retained electrodes FC1 and C2 above the left and right SM1 respectively as those were least affected by the artifact.

Significant ERD in the beta band in each signal selection (left SM1, right SM1, EMG and force) was identified with cluster-based non-parametric permutation tests.^44^ For performing the tests, 1000 surrogate amplitude modulation maps were created from epochs of subband normalized envelopes that were randomly flipped about 1. For both the genuine and each surrogate dataset, we identified the largest cluster of time-frequency bins (bin size, 1 Hz × 0.05 s) exhibiting amplitude suppression exceeding the 95^th^ percentile across all maps. The *p*-value for the genuine cluster size was obtained as the proportion of surrogate cluster size values that were higher than the observed genuine value.

Potential differences in beta ERD cluster size and maximal depth between the SM1, EMG, and force signals were investigated with Bonferroni corrected paired-samples *t*-tests.

The relationship between the size or maximal depth of the significant beta ERD cluster in pairs of signals was assessed with Pearson correlations.

The similarity of the beta ERD between pairs of signals was further evaluated as the Pearson correlation between their time-frequency maps in the window ranging from - 500 ms before to 1500 ms after movement and from 13 to 30 Hz. The significance of this correlation coefficient was assessed with a *t*-test against 0.

#### Movement-synchronized EMG and force amplitude increase

We assessed the extent to which movement execution led to an artifactual increase in EMG and force amplitude, which could have obscured a beta ERD. To this end, we used the same non-parametric permutation test described above to assess the presence and significance of clusters of amplitude increase within amplitude modulation maps in the 5*–*40-Hz frequency range and -200*–*400-ms time-window. We subsequently quantified the area covered by all significant clusters within the sub-beta (4 to 13 Hz), beta (13 to 30 Hz), and supra-beta (30 to 40 Hz) frequency bands. A participant’s EMG or force signals were considered to contain a movement-induced artifact that could have obscured a beta ERD if the artifact-contaminated area in the beta frequency band was larger than 2 Hz s.

#### Interhemispheric and interhand beta envelope coupling

As a preliminary step to estimate interhemispheric beta envelope coupling, MEG signals were projected into the source space. For that, MEG and MRI coordinate systems were co-registered using the 3 anatomical fiducial points for initial estimation and the head-surface points for further manual refinement. Each subject’s MRI was segmented with Freesurfer (Martinos Center for Biomedical Imaging, Boston, MA, RRID:SCR_001847).^45,46^ Then, a non-linear transformation from individual MRIs to the MNI (Montreal Neurological Institute) brain was computed using the spatial normalization algorithm implemented in Statistical Parametric Mapping (SPM8, Wellcome Department of Cognitive Neurology, London, UK, RRID:SCR_007037).^47^ This transformation was used to map a homogeneous 5-mm grid sampling the MNI brain volume onto individual brain volumes. For each subject and grid point, the MEG forward model corresponding to three orthogonal current dipoles was computed using the one-layer Boundary Element Method implemented in the MNE software suite (Martinos Center for Biomedical Imaging, Boston, MA, RRID:SCR_005972). The forward model was then reduced to its first two principal components. This procedure is justified by the insensitivity of MEG to currents radial to the skull, and hence, this dimension reduction leads to considering only the tangential sources. Source signals were then reconstructed with Minimum-Norm Estimates inverse solution.^48^

Source-projected signals were processed as gradiometer pair signals to extract artifact-free segments of subband normalized envelopes. We then estimated the Pearson correlation coefficient between left and right hemisphere envelopes. This correlation analysis was conducted for all pairs of sources located less than 15 mm from previously reported coordinates of the SM1 areas ([*±*42 -26 54] mm), and ensuing correlation values were averaged across all pairs of sources. However, it is known that spatial leakage induces a bias in correlation estimation.^49^ Therefore, a value of correlation corrected for volume leakage was computed from normalized envelopes derived from orthogonalized subband signals. In practice, for each pair of sources, the left-hemisphere subband signal was regressed out from that of the right hemisphere.^49,50^ Since this correction scheme is asymmetric, the corrected correlation was estimated again after inverting the left- and right-hemisphere signals, and both corrected correlation estimates were averaged to provide the final value.

Similarly, interhand envelope correlations were estimated for the EMG signals, and for the force signals. This analysis for the EMG could not be conducted for 2 participants for whom the EMG was lost due to technical issues.

The maximal value of correlation and the corresponding frequency were extracted across the 13–30 Hz frequency range, and for MEG, also across all pairs of sources. The relationships between interhemispheric and interhand coupling metrics (correlation and peak frequency) were assessed with Pearson correlations.

## Data Availability

Preprocessed data from each task has been deposited on the Open Science Framework (OSF; https://osf.io/9fgw8/overview). All custom Matlab and R code used for the data preprocessing, analyses, and visualization is also made publicly available on the OSF.

## Acknowledgements

**C.G.** was supported by an Aspirant Research Fellowship awarded by the Fonds de la Recherche Scientifique (FNRS; Brussels, Belgium; grant 1.A.211.24F).

**P.C.** was supported by a Clinical Researcher Fellowship awarded by the F.R.S.-FNRS (Brussels, Belgium; grant 40024164). **S.J.M.** was supported by an Aspirant Research Fellowship awarded by the FNRS (Brussels, Belgium; grant FC 46249). **X.D.T.** is Clinical Researcher at the FNRS. **G.N.** is a postdoctorate Clinical Master Specialist at the FNRS (Brussels, Belgium). **M.B., N.Y-M.,** and **D.D.** were supported by the Brussels-Wallonia Federation (Collective Research Initiatives grant) and the Walloon Region (FRFS-WELBIO strategic axis).

The MEG project at the Université libre de Bruxelles and Hôpital Universitaire de Bruxelles is financially supported by the Fonds Erasme (Brussels, Belgium; Research Convention “Les Voies du Savoir”).

The PET-MR project at the Université libre de Bruxelles and Hôpital Universitaire de Bruxelles is financially supported by the Association Vinçotte Nuclear (AVN, Brussels, Belgium).

## Author contributions

**C.G.** acquired the data, performed data analyses and data visualisations, and wrote the first draft of the manuscript; **P.C., N.Y-M.,** and **D.D.** acquired the data and performed data analyses; **S.J.M., A.H.M.T., and L.M.** assisted the participant recruitment and data collection; **V.W.** assisted with MEG source reconstruction and connectivity analyses; **G.N.** and **X.D.T.** assisted with project conceptualisation and supervision; **M.B.** conceived the project, obtained the funding, developed the analysis software, and supervised the data collection, analysis, and visualization. All authors participated in result interpretation and provided comments on the final draft of the manuscript.

## References

1. Buzsáki, G. & Draguhn, A. Neuronal oscillations in cortical networks. Science 304, 1926–1929 (2004).

2. Beste, C., Münchau, A. & Frings, C. Towards a systematization of brain oscillatory activity in actions. Commun Biol 6, 137 (2023).

3. Berger, H. Das Elektrenkephalogramm des Menschen. (1933).

4. Heinrichs-Graham, E. et al. Neuromagnetic evidence of abnormal movement-related beta desynchronization in Parkinson’s disease. Cereb Cortex 24, 2669–2678 (2014).

5. Espenhahn, S. et al. Sensorimotor cortex beta oscillations reflect motor skill learning ability after stroke. Brain Commun 2, fcaa161 (2020).

6. Buard, I., Kronberg, E., Steinmetz, S., Hepburn, S. & Rojas, D. C. Neuromagnetic Beta-Band Oscillations during Motor Imitation in Youth with Autism. Autism Res Treat 2018, 9035793 (2018).

7. Moran, J. K. & Senkowski, D. Intersensory attention deficits in schizophrenia relate to ongoing sensorimotor beta oscillations. Schizophrenia (Heidelb) 11, 19 (2025).

8. Peter, J. et al. Movement-related beta ERD and ERS abnormalities in neuropsychiatric disorders. Front Neurosci 16, 1045715 (2022).

9. Bonaiuto, J. J. et al. Laminar dynamics of high amplitude beta bursts in human motor cortex. Neuroimage 242, 118479 (2021).

10. Shin, H., Law, R., Tsutsui, S., Moore, C. I. & Jones, S. R. The rate of transient beta frequency events predicts behavior across tasks and species. Elife 6, (2017).

11. Szul, M. J. et al. Diverse beta burst waveform motifs characterize movement-related cortical dynamics. Prog Neurobiol 228, 102490 (2023).

12. Engel, A. K. & Fries, P. Beta-band oscillations--signalling the status quo? Curr. Opin. Neurobiol. 20, 156–165 (2010).

13. Démas, J. et al. Mu rhythm: State of the art with special focus on cerebral palsy. Ann. Phys. Rehabil. Med. 63, 439–446 (2020).

14. Kilavik, B. E., Zaepffel, M., Brovelli, A., MacKay, W. A. & Riehle, A. The ups and downs of beta oscillations in sensorimotor cortex. Experimental Neurology vol. 245 15–26 Preprint at 10.1016/j.expneurol.2012.09.014 (2013).

15. Diesburg, D. A., Greenlee, J. D. & Wessel, J. R. Cortico-subcortical β burst dynamics underlying movement cancellation in humans. Elife 10, (2021).

16. Pfurtscheller, G. & Lopes da Silva, F. H. Event-related EEG/MEG synchronization and desynchronization: basic principles. Clin. Neurophysiol. 110, 1842–1857 (1999).

17. Enz, N., Ruddy, K. L., Rueda-Delgado, L. M. & Whelan, R. Volume of β-Bursts, But Not Their Rate, Predicts Successful Response Inhibition. J Neurosci 41, 5069–5079 (2021).

18. Zrenner, C., Desideri, D., Belardinelli, P. & Ziemann, U. Real-time EEG-defined excitability states determine efficacy of TMS-induced plasticity in human motor cortex. Brain Stimul. 11, 374–389 (2018).

19. Salenius, S. & Hari, R. Synchronous cortical oscillatory activity during motor action. Curr Opin Neurobiol 13, 678–684 (2003).

20. Mary, A. et al. Age-related differences in practice-dependent resting-state functional connectivity related to motor sequence learning. Hum. Brain Mapp. 38, 923–937 (2017).

21. Sugata, H. et al. Role of beta-band resting-state functional connectivity as a predictor of motor learning ability. Neuroimage 210, 116562 (2020).

22. Mongold, S. J. et al. Temporally stable beta sensorimotor oscillations and corticomuscular coupling underlie force steadiness. Neuroimage 261, 119491 (2022).

23. Baker, S. N., Pinches, E. M. & Lemon, R. N. Synchronization in monkey motor cortex during a precision grip task. II. effect of oscillatory activity on corticospinal output. J. Neurophysiol. 89, 1941–1953 (2003).

24. Bräcklein, M., Barsakcioglu, D. Y., Del Vecchio, A., Ibáñez, J. & Farina, D. Reading and Modulating Cortical β Bursts from Motor Unit Spiking Activity. J. Neurosci. 42, 3611–3621 (2022).

25. Echeverria-Altuna, I. et al. Transient beta activity and cortico-muscular connectivity during sustained motor behaviour. Prog Neurobiol 214, 102281 (2022).

26. Georgiev, C. et al. Transcallosal generation of phase-aligned beta bursts underlies TMS-induced interhemispheric inhibition. Imaging Neurosci (Camb) 3, imag_a_00570 (2025).

27. Bourguignon, M. et al. MEG Insight into the Spectral Dynamics Underlying Steady Isometric Muscle Contraction. J Neurosci 37 10421–10437 (2017).

28. Johnson, A. N., & Shinohara, M. Corticomuscular coherence with and without additional task in the elderly. J Appl Physiol 112, 970–981 (2012).

29. Bourguignon, M., Jousmäki, V., Dalal, S. S., Jerbi, K. & De Tiège, X. Coupling between human brain activity and body movements: Insights from non-invasive electromagnetic recordings. Neuroimage 203, 116177 (2019).

30. Barone, J. & Rossiter, H. E. Understanding the Role of Sensorimotor Beta Oscillations. Front. Syst. Neurosci. 15, 655886 (2021).

31. Barry, R. J. & De Blasio, F. M. EEG differences between eyes-closed and eyes-open resting remain in healthy ageing. Biol Psychol 129, 293–304 (2017).

32. Rempe, M. P. et al. Spontaneous sensorimotor beta power and cortical thickness uniquely predict motor function in healthy aging. Neuroimage 263, 119651 (2022).

33. Georgiev, C. et al. Gaining insight into the nonfocality of beta oscillation suppression along the sensorimotor cortex using corticomuscular coherence. Eur J Neurosci 63, e70600 (2026).

34. Cordier, A. et al. The dissociative role of bursting and non-bursting neural activity in the oscillatory nature of functional brain networks. Imaging Neurosci (Camb) 2, imag-2-00231 (2024).

35. Hari, R. et al. Human primary motor cortex is both activated and stabilized during observation of other person’s phasic motor actions. Philos. Trans. R. Soc. Lond. B Biol. Sci. 369, 20130171 (2014).

36. Pavlidou, A., Schnitzler, A. & Lange, J. Beta oscillations and their functional role in movement perception. Transl. Neurosci. 5, 286–292 (2014).

37. van Wijk, B. C. M., Beek, P. J. & Daffertshofer, A. Differential modulations of ipsilateral and contralateral beta (de)synchronization during unimanual force production. Eur J Neurosci 36, 2088–2097 (2012).

38. Piitulainen, H. et al. Phasic stabilization of motor output after auditory and visual distractors. Hum Brain Mapp 36, 5168–5182 (2015).

39. Oldfield, R. C. The assessment and analysis of handedness: The Edinburgh inventory. Neuropsychologia 9 97–113 (1971).

40. Taulu, S. & Simola, J. Spatiotemporal signal space separation method for rejecting nearby interference in MEG measurements. Phys Med Biol 51, 1759–1768 (2006).

41. Vigário, R., Särelä, J., Jousmäki, V., Hämäläinen, M. & Oja, E. Independent component approach to the analysis of EEG and MEG recordings. IEEE Trans. Biomed. Eng. 47, 589–593 (2000).

42. Hyvärinen, A. & Oja, E. Independent component analysis: algorithms and applications. Neural Netw 13, 411–430 (2000).

43. Bigdely-Shamlo, N., Mullen, T., Kothe, C., Su, K.-M. & Robbins, K. A. The PREP pipeline: standardized preprocessing for large-scale EEG analysis. Front. Neuroinform. 9, 16 (2015).

44. Maris, E. & Oostenveld, R. Nonparametric statistical testing of EEG- and MEG-data. J. Neurosci. Methods 164, 177–190 (2007).

45. Reuter, M., Schmansky, N. J., Rosas, H. D. & Fischl, B. Within-subject template estimation for unbiased longitudinal image analysis. Neuroimage 61, 1402–1418 (2012).

46. Fischl, B. FreeSurfer. Neuroimage 62, 774–781 (2012).

47. Ashburner, J. & Friston, K. J. Nonlinear spatial normalization using basis functions. Hum Brain Mapp 7, 254–266 (1999).

48. Bertels, J. et al. Neurodevelopmental oscillatory basis of speech processing in noise. Dev Cogn Neurosci 59, 101181 (2023).

49. Wens, V. et al. A geometric correction scheme for spatial leakage effects in MEG/EEG seed-based functional connectivity mapping. Hum Brain Mapp 36, 4604–4621 (2015).

50. Brookes, M. J., Woolrich, M. W. & Barnes, G. R. Measuring functional connectivity in MEG: a multivariate approach insensitive to linear source leakage. Neuroimage 63, 910–920 (2012).

