## Supplementary Material for "Assessment of sensorimotor cortical beta oscillations from peripheral electromyography and force recordings"

#### Table of contents

S1. Beta ERS 2

S2. Beta Bursts 3

S3. Burstiness of $MEG_{SM1}$ and peripheral signals 4

Supplementary Fig. 1 6

Supplementary Fig. 2 8

Supplementary Fig. 3 9

Supplementary Fig. 4 10

Supplementary Table 1 11

Supplementary Table 2 12

Supplementary Table 3 13

Supplementary Table 4 14

Supplementary Table 5 15

References 16

**S1. Beta ERS**

Having mostly circumvented the issue of unstable EMG and force during the movement allowed us to use the data from the *contralateral upper limb movement execution task* to also investigate the post-movement beta ERS. These results revealed that the left finger movement led to a beta ERS in the left and right $MEG_{SM1}$, along with a beta ERS in the right EMG and force signals in the time interval from 0.5 to 2.5 s (Supplementary Table 2). The beta ERS size was significantly smaller in the EMG ($t_{40}$ = 2.74, $p_{corrected}$ = 0.018) and the force signal ($t_{40}$ = 3.68, $p_{corrected}$ = 0.001) compared to the left $MEG_{SM1}$. There was no significant difference in ERS peak magnitude between the left $MEG_{SM1}$ and the right EMG ($t_{40}$ = -0.14, $p_{uncorrected}$ = 0.89), however, ERS peak magnitude was significantly higher in the left $MEG_{SM1}$ compared to the right force signal ($t_{40}$ = 4.66, $p_{corrected}$ < 0.001). A cross-modality correlation analysis revealed no significant correlations between left $MEG_{SM1}$ and EMG beta ERS parameters (size, $r$ = 0.30, $p$ = 0.059; peak magnitude, $r$ = -0.06, $p$ = 0.72) and a significant positive correlation between left $MEG_{SM1}$ and force beta ERS parameters (size, $r$ = 0.56, $p$ < 0.001; peak magnitude, $r$ = 0.64, $p$ < 0.001).

In the validation recordings, a significant beta ERS in the time interval from 0.5 to 2.5 s relative to left finger movement was identified in 11 (52%) of the young participants, and in 10 (77%) of the older participants in the force signal (all $p$ < 0.05). The beta ERS was greater in older compared to young participants in terms of size (young, 1.6 ± 1.8 Hz s; older, 4.8 ± 5.5 Hz s; $t_{32}$ = -2.45, $p$ = 0.020), however, there were no significant differences in peak magnitude (young, 12.2 ± 7.5%; older, 17.7 ± 10.7%; $t_{32}$ = -1.75, $p$ = 0.090).

In patients with Parkinson's disease, a significant beta ERS in the time interval from 0.5 to 2.5 s was identified in 13 (100%) patients for both the left and right $EEG_{SM1}$ and in 12 (92%) patients for the force (all $p$ < 0.05; Supplementary Table 5). However, as reported in the main text, the identified force beta ERS in 2 patients was likely a movement-synchronized amplitude increase. Beta ERS parameters were not significantly different between the $EEG_{SM1}$ and the force (size, $t_{10}$ = -1.15, $p$ = 0.28; peak magnitude, $t_{10}$ = 1.07, $p$ = 0.31). These results were not affected by the exclusion of the two participants with a movement-synchronized force amplitude increase.

**S2. Beta Bursts**

We investigated how the observed beta ERD in the $MEG_{SM1}$, EMG, and force signals during contralateral finger movement related to the bursty nature of the beta oscillations. For that, we assessed the individual beta bursts detected in each signal relative to movement onset, for each participant separately. To obtain beta envelopes, the 9 $MEG_{SM1}$ signals, the EMG signal, and the force signal were filtered between 13 and 30 Hz, the filtered $MEG_{SM1}$ signals were reduced to their first principal component, and the signals' envelope was extracted with Hilbert transformation. The envelopes were further normalized by their <0.1 Hz trend. Then, for each signal and for each participant, time intervals where the envelope exceeded its 75$^{th}$ percentile were identified and considered as beta bursts if their duration was longer than 50 ms.[1] Based on this, for each movement onset, we extracted the probability of beta burst occurrence in 3 ERD time windows: -100 to 100 ms, 100 to 300 ms, and 300 to 500 ms relative to finger movement onset. Then, the probability in each time window was compared against baseline values obtained in the time intervals from -900 to -500 ms and from 1100 to 2300 ms with a paired *t*-test across movements with false discovery rate correction. Likewise, we compared the duration of all the bursts detected within each of the 3 time windows to that of the bursts detected during the baseline interval with a two-sample *t*-test with false discovery rate correction. In this setup, we compared the extent to which beta bursts were suppressed in the time in which a beta ERD is expected to occur.

The left finger movement execution resulted in a significant reduction in the probability of beta burst occurrence in the left $MEG_{SM1}$ in at least one ERD time window in 27 (66%) participants. Similar modulations were observed in the right EMG in 22 (54%) participants and in the force in 25 (61%) participants. The mean probability of beta burst occurrence across the 3 ERD time windows was 31 ± 1% in the left $MEG_{SM1}$, 41 ± 14% in the right EMG, and 36 ± 12% in the force.

In addition, the left finger movement execution resulted in a significant reduction in beta burst duration in the left $MEG_{SM1}$ in at least one ERD time window in 9 (22%) participants. Similar modulations were observed in the right EMG in 14 (34%) participants and in the force signal in 16 (39%) participants. The mean beta burst duration across the 3 ERD time windows was 103 ± 12 ms in the left $MEG_{SM1}$, 110 ± 27 ms in the right EMG, and 114 ± 18 ms in the force.

#### S3. Burstiness of $MEG_{SM1}$ and peripheral signals

To confirm that interhemispheric and interhand beta envelope couplings during the bimanual contraction task depended on the presence of beta bursts, we assessed the association between the magnitude of these couplings and a measure of the salience of these bursts, termed burstiness.[2] The burstiness of MEG gradiometer signals, source-reconstructed left- and right SM1, rFDI EMG and force was computed as the coefficient of variation of their subband envelopes,[2] which were centered on 5–40 Hz and obtained as explained in the methods (subsection ***"Data Analysis"***). The burstiness takes value ~0.483 for a white noise, and increases when bursts are present.[2]

We estimated beta-band burstiness as the maximum burstiness value within 15–25 Hz, and across the left or right $MEG_{SM1}$ sensor selection for $MEG_{SM1}$, or across the left- or right source-selections for source-reconstructed SM1 signals. Group-level differences between each signal's beta-band burstiness and that of a white noise were investigated with one-sample *t*-tests. In this comparison, values for the white noise were also estimated as the maximum across the beta band for peripheral signals (0.494 on average) and across the beta band for the 9 signals for $MEG_{SM1}$ (0.502 on average). Associations between beta-band couplings (interhemispheric or interhand) and signals' beta-band burstiness were investigated with Pearson correlations. In this analysis, beta-band burstiness was averaged across the two brain hemispheres or across the two hands. Lastly, association between each SM1's beta-band burstiness and that of each contralateral peripheral signal was investigated with Pearson correlation across participants.

In line with previous reports,[2] $MEG_{SM1}$ and peripheral signals' burstiness peaked in the beta band, and, for MEG, above bilateral SM1 hand areas (Supplementary Fig. 1a). In addition, all signals' burstiness was significantly greater than that of corresponding white noise (all $p < 0.001$; Supplementary Table 3; Supplementary Fig. 1a), therefore demonstrating the presence of beta bursts in each $MEG_{SM1}$ and peripheral signal.

Similarly, in line with previous reports,[3] there was a strong positive correlation between the source-reconstructed bilateral $MEG_{SM1}$ beta envelope coupling and the pooled $MEG_{SM1}$ burstiness ($r$ = 0.95, $p$ < 0.0001; Supplementary Fig. 1b), indicating that stronger interhemispheric coupling coincided with stronger burstiness across hemispheres. Likewise, we found a significant positive correlation between beta interhand coupling and the pooled burstiness for the EMG signals ($r$ = 0.76, $p$ < 0.0001) and a significant positive correlation for the force signals ($r$ = 0.83, $p$ < 0.0001; Supplementary Fig. 1b). These results show that interhand beta envelope couplings are driven by the presence of beta bursts in peripheral signals, as is the case at the cortical level.

Further supporting the notion that peripheral beta bursts originate from SM1 beta bursts, we found significant positive correlations between the left $MEG_{SM1}$ beta-band burstiness and that of the right EMG and force (Supplementary Fig. 1c). A significant positive correlation was also found between the right $MEG_{SM1}$ beta-band burstiness and that of the left EMG and force (Supplementary Fig. 1c).

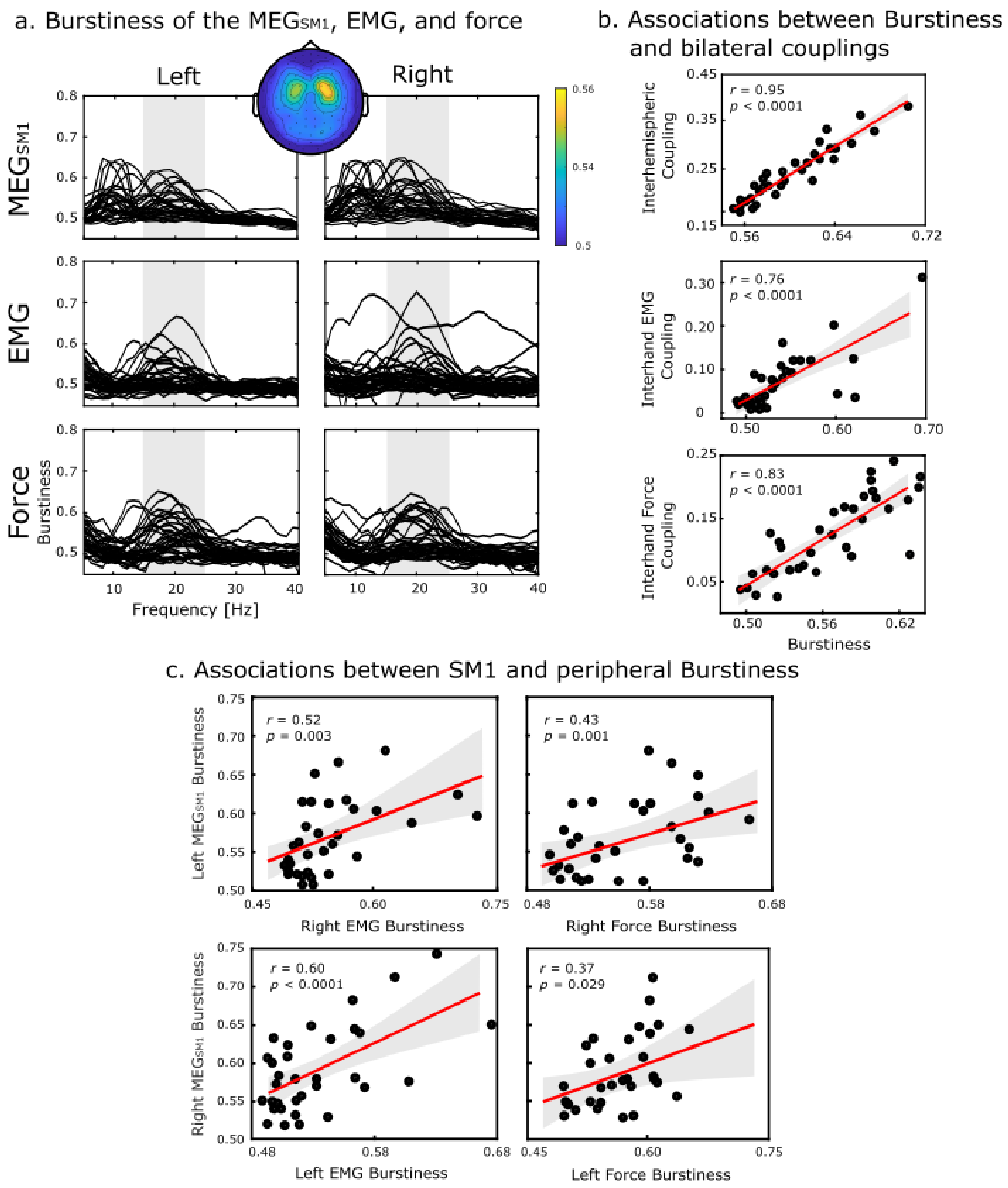


*Supplementary Fig. 1. Results of the burstiness analysis.* ***a.*** *Burstiness of each $MEG_{SM1}$, EMG, and force signal during the bimanual contraction task as function of the frequency. The topographic map presents the distribution of the maximum*

*burstiness within the 15–25 Hz range.* ***b.*** *Pearson correlations between beta-band envelope couplings and burstiness. This relation is presented for interhemispheric* $MEG_{SM1}$ *coupling and SM1 burstiness, interhand EMG coupling and EMG burstiness, and interhand force coupling and force burstiness. Regression lines are in red and associated confidence areas are in gray.* ***c.*** *Pearson correlations between pooled* $MEG_{SM1}$ *burstiness and the contralateral EMG and force burstiness. Regression lines are in red and associated 95% confidence areas are in gray.*

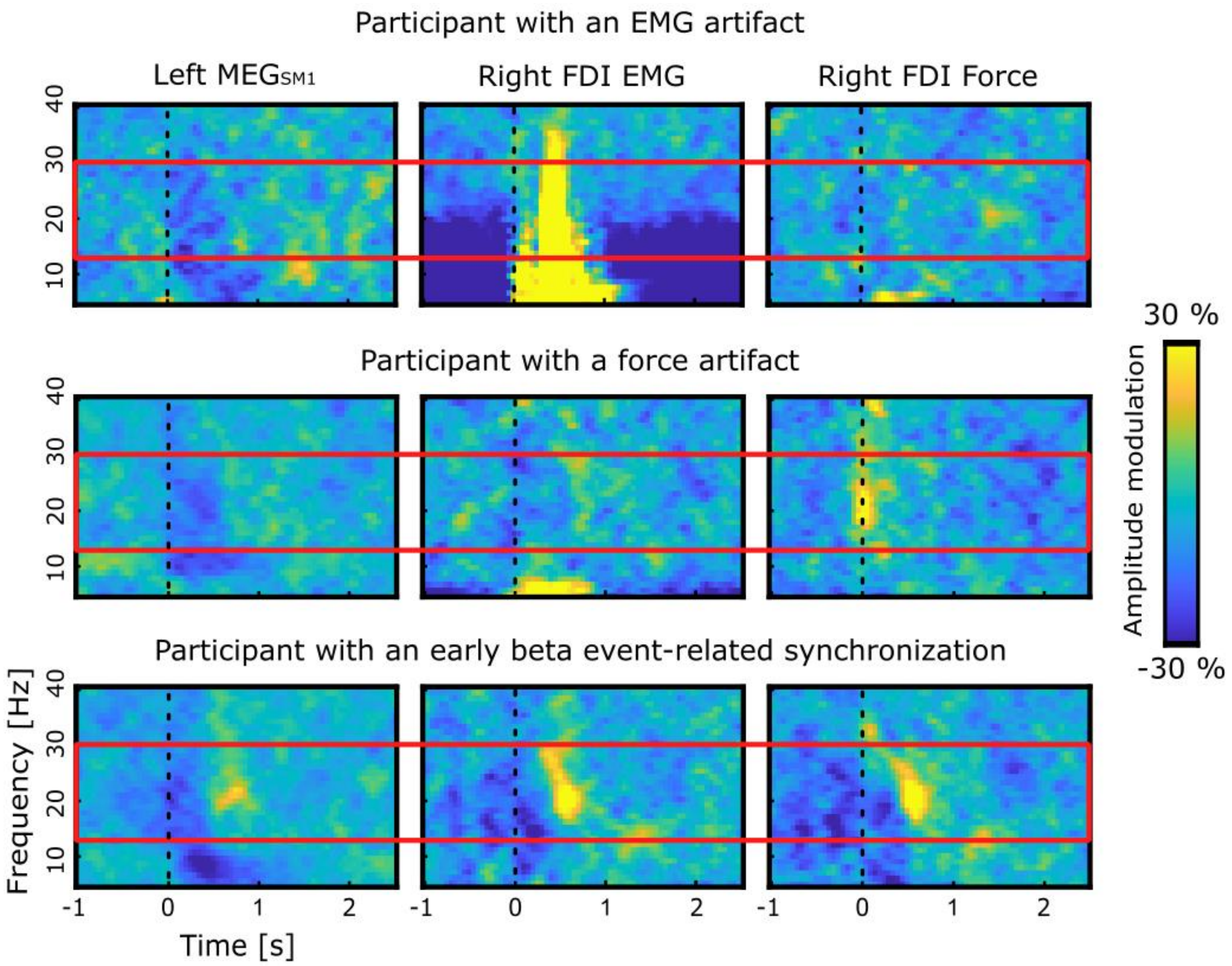


*Supplementary Fig. 2. Time-frequency amplitude modulation of left $MEG_{SM1}$, right FDI EMG, and right hand force signal for three participants with task-induced amplitude increase in the peripheral signals in the contralateral upper limb movement execution task. The figure demonstrates a participant with a significant movement-synchronized amplitude increase in the EMG (row 1), a participant with a significant movement-synchronized amplitude increase in the force (row 2), and the participant with a task-induced increase in both peripheral signals consistent with a post-movement beta event-related synchronization (row 3). Dotted black lines at time 0 indicate movement onset and red rectangles outline the classical beta frequency range (13–30 Hz).*

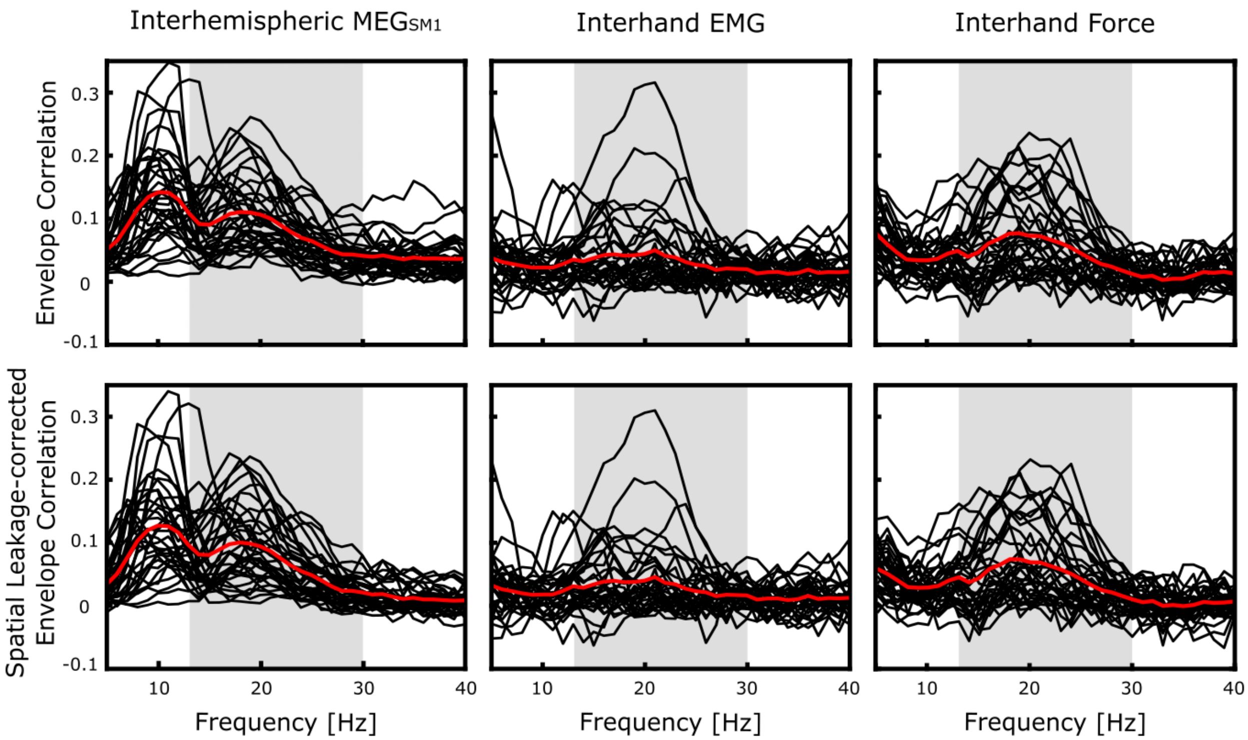


*Supplementary Fig. 3. Spectra of interhemispheric and interhand envelope correlations, without (top) and with signal orthogonalization (bottom). Black lines represent single participant values and red lines represent mean values across participants. Gray shaded areas outline the classical beta frequency range (13–30 Hz).*

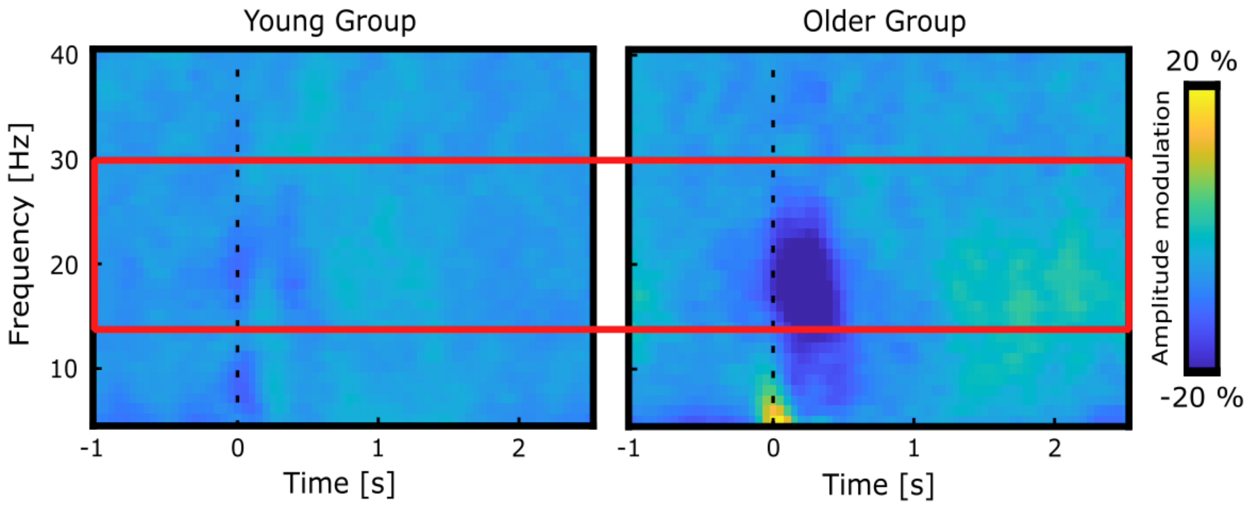


*Supplementary Fig. 4. Grand averaged time-frequency amplitude modulation in the right force signals of the young (left) and older (right) group in the validation contralateral movement execution experiment in standard laboratory settings. Dotted black lines at time 0 indicate movement onset and red rectangles outline the classical beta frequency range (13–30 Hz).*

| | Ipsilateral Upper Limb Movement Execution Task | | |
|---|---|---|---|
| | **ERD Cluster Size [Hz s]** | **ERD Cluster Depth [%]** | **ERD Significance [N]** |
| **Right $MEG_{SM1}$** | 3.1 ± 2.4 | 31.0 ± 5.9 | 10 (71%) |
| **Left $MEG_{SM1}$** | 7.6 ± 5.5 | 37.9 ± 9.7 | 14 (100%) |
| **Right EMG** | 2.5 ± 3.0 | 33.9 ± 11.2 | 9 (64%) |
| **Right Force** | 2.9 ± 2.4 | 32.5 ± 7.3 | 14 (100%) |

*Supplementary Table 1* **|** *Beta ERD cluster size, depth and statistical significance ($p < 0.05$) for the $MEG_{SM1}$, the right FDI EMG, and the right force in the ipsilateral upper limb movement execution tasks. Data are presented as mean ± SD or number of participants (%).*

| | Contralateral Upper Limb Movement Execution Task | | | | | |
|---|---|---|---|---|---|---|
| | **ERD Cluster Size [Hz s]** | **ERD Cluster Depth [%]** | **ERD Significance [N]** | **ERS Cluster Size [Hz s]** | **ERS Cluster Peak [%]** | **ERS Significance [N]** |
| **Right $MEG_{SM1}$** | 8.0 ± 5.5 | 37.6 ± 6.3 | 37 (90%) | 5.2 ± 4.9 | 30.4 ± 9.6 | 34 (83%) |
| **Left $MEG_{SM1}$** | 6.4 ± 4.5 | 37.2 ± 7.5 | 40 (98%) | 2.8 ± 3.0 | 26.5 ± 8.3 | 25 (61%) |
| **Right EMG** | 2.1 ± 2.8 | 30.9 ± 10.2 | 29 (71%) | 1.4 ± 2.3 | 27.3 ± 32.8 | 14 (34%) |
| **Right Force** | 2.4 ± 3.3 | 31.6 ± 7.7 | 25 (61%) | 1.4 ± 1.7 | 21.1 ± 9.1 | 21 (51%) |

*Supplementary Table 2* | *Beta ERD cluster size, depth, and statistical significance ($p < 0.05$) as well as ERS cluster size, peak magnitude, and statistical significance ($p < 0.05$) for the $MEG_{SM1}$, the right FDI EMG, and the right force in the contralateral upper limb movement execution tasks. Data are presented as mean ± SD or number of participants (%).*

| | Burstiness | |
|---|---|---|
| | **Left** | **Right** |
| **MEG$_{SM1}$** | 0.57 ± 0.05 ******* | 0.59 ± 0.06 ******* |
| | **Right** | **Left** |
| **EMG** | 0.55 ± 0.06 ******* | 0.53 ± 0.04*** |
| **Force** | 0.56 ± 0.04 ******* | 0.57 ± 0.05 ******* |

*Supplementary Table 3* **|** *Burstiness of each SM1 and peripheral signal during the bimanual contraction task. Data are presented as mean ± SD, with the significance in comparison to values derived from a white noise indicated aside.*** = p < 0.001.*

| | |
|---|---|
| **Age** | 67 ± 6 years |
| **Male** | 6 (46%) |
| **Right-handed** | 9 (69%) |
| **Disease Duration** | 10 ± 6 years |
| **Type** | |
| • **Akineto-rigid dominant subtype** | 8 (62%) |
| • **Tremor dominant subtype** | 5 (38%) |
| **Hoehn and Yahr score** | 2 ± 0 |
| **MDS-UPDRS III** | 29 ± 12 |
| • **Akineto-rigid subscore** | 17 ± 7 |
| • **Tremor subscore** | 6 ± 4 |

*Supplementary Table 4* | *Descriptive statistics of Parkinson's disease patients. Data are presented as mean ± SD or number of participants (%). MDS-UPDRS, Movement Disorders Society Unified Parkinson's Disease Rating Scale.*

| | Beta ERD | | Beta ERS | |
|---|---|---|---|---|
| | **ERD Cluster Size [Hz s]** | **ERD Cluster Depth [%]** | **ERS Cluster Size [Hz s]** | **ERS Cluster Peak [%]** |
| **Right $EEG_{SM1}$** | 11.7 ± 3.9 | 37.2 ± 7.8 | 8.3 ± 5.3 | 25.4 ± 7.8 |
| **Left $EEG_{SM1}$** | 10.5 ± 4.2 | 36.0 ± 8.0 | 6.5 ± 4.4 | 23.1 ± 7.3 |
| **Force** | 7.1 ± 3.5 | 32.0 ± 8.3 | 9.7 ± 6.5 | 20.5 ± 9.9 |

*Supplementary Table 5* **|** *$EEG_{SM1}$ and force beta ERD cluster size and depth as well as ERS cluster size and peak from the Parkinson's disease task. Data are presented as mean ± SD.*

### References

1. Georgiev, C. *et al.* Transcallosal generation of phase-aligned beta bursts underlies TMS-induced interhemispheric inhibition. *Imaging Neurosci (Camb)* vol. 3, imag_a_00570 (2025). doi:10.1162/imag_a_00570

2. Bourguignon, M. *et al.* MEG Insight into the Spectral Dynamics Underlying Steady Isometric Muscle Contraction. *The Journal of Neuroscience* vol. 37, 43 (2017): 10421-10437. doi:10.1523/JNEUROSCI.0447-17.2017

3. Cordier, A. *et al.* The dissociative role of bursting and non-bursting neural activity in the oscillatory nature of functional brain networks. *Imaging Neurosci (Camb)* vol. 2, imag-2-00231 (2024). doi:10.1162/imag_a_00231
